# Duration of Morning Hyperinsulinemia Determines Hepatic Glucose Uptake and Glycogen Storage Later in the Day

**DOI:** 10.1101/2024.05.10.593551

**Authors:** Hannah L. Waterman, Mary Courtney Moore, Marta S. Smith, Ben Farmer, Melanie Scott, Dale S. Edgerton, Alan D. Cherrington

## Abstract

The second meal phenomenon refers to the improvement in glucose tolerance seen following a second identical meal. We previously showed that 4 hours of morning hyperinsulinemia, but not hyperglycemia, enhanced hepatic glucose uptake (HGU) and glycogen storage during an afternoon hyperinsulinemic-hyperglycemic (HIHG) clamp. Our current aim was to determine if the duration or pattern of morning hyperinsulinemia is important for the afternoon response to a HIHG clamp. To determine this, we administered the same total amount of insulin either over 2h in the first (Ins2h-A) or second (Ins2h-B) half of the morning, or over the entire 4h (Ins4h) of the morning. In the 4h afternoon period, all three groups had 4x-basal insulin, 2x-basal glycemia, and portal glucose infusion to expose the liver to the primary postprandial regulators of hepatic glucose metabolism. During the afternoon clamp, there was a marked increase in HGU and hepatic glycogen synthesis in the Ins4h group compared to the Ins2h-A and Ins2h-B groups, despite matched hepatic glucose loads and total insulin infusion rates. Thus, the longer duration (Ins4h) of lower hyperinsulinemia in the morning seems to be the key to much greater liver glucose uptake during the afternoon clamp.

**New and noteworthy:** Morning insulin exposure primes the liver for increased hepatic glucose uptake and glycogen storage during a subsequent hyperinsulinemic-hyperglycemic clamp. This study addressed whether the pattern and/or duration of insulin delivery in the morning influences insulin’s ensuing priming effect. We found that despite receiving equal total doses of insulin in the morning, a prolonged, lower rate of morning insulin delivery improved afternoon liver glucose metabolism more effectively than a shorter, higher rate of delivery.

## Introduction

Individuals who regularly skip breakfast have an increased risk of becoming obese, developing type 2 diabetes, and being diagnosed with cardiovascular disease (1–7). While there is an association between breakfast consumption and improved metabolic health, the nutritional composition of the first meal of the day is an important factor to consider when assessing the positive health benefits associated with breakfast (8–11). Specifically, it has been shown that people who consume breakfasts rich in fibrous whole grains, low-fat dairy, nuts, and fruit display a lower BMI and amount of abdominal fat compared to those who consume breakfast meals composed of high-fat dairy products, processed grains, and artificial sugars (12–14). Moreover, dietary patterns that are high in rapidly available carbohydrates rather than slower absorbing carbohydrates are correlated with less favorable metabolic outcomes (15, 16).

One of the primary responses to a carbohydrate-rich meal is the stimulation of insulin secretion, which acts to bring circulating glucose back to basal levels. Meals that are rich in simple carbohydrates, such as refined sugars and processed foods, can lead to a short-lived episode of hyperinsulinemia. Both insulin and glucose levels begin to rise immediately after consuming rapidly absorbed carbohydrates, reaching a peak within 30 minutes to an hour, and remain elevated for only two hours after consumption (17, 18). In contrast, meals that are higher in complex carbohydrates tend to result in a more prolonged, lower bout of hyperinsulinemia (19–22). Furthermore, the level of hyperinsulinemia is substantially greater after absorbing simple carbohydrates versus complex carbohydrates. For example, one study showed that raw starch ingestion resulted in 35-65% lower insulin response when compared to glucose or sucrose ingestion (23). This glycemic response is worse in individuals with type 2 diabetes due to an insufficient secretion of insulin following ingestion of a carbohydrate, leading to abnormally high postprandial glucose levels (24–26). Thus, it is possible to influence the duration and extent of morning hyperinsulinemia by adjusting meal composition and size.

Human studies have shown that despite being fed an identical meal at breakfast and lunch, glucose and insulin excursions were markedly reduced in response to a second meal. This phenomenon, otherwise known as the “second meal phenomenon” or “Staub-Traugott effect”, has been observed in both healthy individuals and those with diabetes, although some studies suggest that it is abnormal in the latter (27–31). The underlying mechanism explaining this response has yet to be fully uncovered. In previous studies, our lab showed that morning hyperinsulinemia per se is a critical component in priming the liver to increase HGU and glycogen storage later in the day (32). It was not clear, however, how the duration or kinetics of the rise in morning insulin influenced the priming effect. These questions are addressed in the current report and have the potential to be particularly important for individuals with diabetes, as those who utilize insulin therapy for diabetes management face an elevated risk of experiencing bouts of low blood sugar due to discrepancies in insulin delivery (33, 34). Thus, learning how to maximize the morning priming effect using lifestyle interventions, such as changing meal composition and/or adjusting the amount of insulin given with each meal, may prove to be an effective way to improve blood sugar regulation.

## Materials and Methods

### Animal care and surgical procedures

Experiments were conducted on both male and female adult mongrel dogs (22.3 ± 0.6 kg) purchased from a USDA-licensed vendor and housed and cared for according to the standards published by the American Association for the Accreditation of Laboratory Animal Care. No sex differences were detected. The dogs were fed a chow and meat diet (46% carbohydrate, 34% protein, 14.5% fat, and 5.5% fiber). Approximately 2 weeks before being studied, each dog underwent a laparotomy during which catheters were inserted into the left common hepatic vein, the hepatic portal vein, and the femoral artery, and infusion catheters into the splenic vein, jejunal vein, and inferior vena cava, and buried in a subcutaneous pocket (35, 36). Ultrasonic blood flow probes were placed around the hepatic artery and the portal vein. The experimental protocol was approved by the Vanderbilt IACUC. All dogs were fasted for 18 hours prior to being studied. Only healthy dogs that consumed at least 75% of their last meal, had a leukocyte count <18,000 mm^3^, and a hematocrit >34 were studied. The total volume of blood withdrawn was <20% of the animal’s blood volume and there was no significant decrease in hematocrit during any of the studies.

### Experimental Design

#### Morning (AM) Clamp

The experimental protocol consisted of two clamp periods **(Fig. 1).** At the beginning of each experiment, the catheters and flow probes were removed from their subcutaneous pockets under local anesthesia. Animals were allowed to rest in a harness for the remainder of the experimental period. Angiocaths were inserted into the cephalic veins for peripheral infusions. Our previous study revealed that it was the elevation of insulin in the morning, not glucose, that was critical for priming the liver for increased glucose uptake and storage (32). Therefore, to generate the insulin priming effect, the animals underwent an AM hyperinsulinemic-euglycemic clamp from 0-240 min, 0-120 min, or 120-240 min (Ins4h, Ins2h-A, or Ins2h-B; *n*=6/group). Thus, the AM clamp began at the same time for the Ins4h group and the Ins2h-A group, and 2 hours later for the Ins2h-B group **(Fig. 1)**. Somatostatin was infused into the inferior vena cava (0.8 µg/kg/min; Bachem, Torrance, CA) to inhibit endogenous pancreatic hormone secretion. Glucagon was replaced intraportally (0.57 ng/kg/min; GlucaGen, Boehringer Ingelheim, Ridgefield, CT) to maintain its basal levels. The insulin (Novolin R; Novo Nordisk, Basværd, Denmark) infusion rates used in the Ins4h group were selected to mimic the rise in endogenous insulin secretion previously observed during a 4h AM duodenal glucose infusion (2.1 mU/kg/min [0-30 min], 2.4 mU/kg/min [30-60 min], and 1.5 mU/kg/min [60-240 min]) (32, 37). These rates were doubled and infused into the hepatic portal vein over 2h (4.2 mU/kg/min [0-15 and 120-135 min], 4.8 mU/kg/min [15-30 and 135-150 min], and 3.0 mU/kg/min [30-120 and 150-240 min]) in the Ins2h-A and Ins2h-B groups, respectively, to match the total amount of insulin being infused in the 4h group (405 mU/kg total insulin over the AM clamp period). Plasma insulin levels remain within a physiologic range with these rates (38, 39). 50% dextrose was infused into a leg vein as needed to maintain euglycemia. Heparinized saline was infused into the femoral artery catheter during the study to replace fluids. At the end of the AM clamp, all hormone infusions ended. Glucose was infused into the inferior vena cava as necessary until the animals were able to maintain euglycemia.

**Figure 1.**
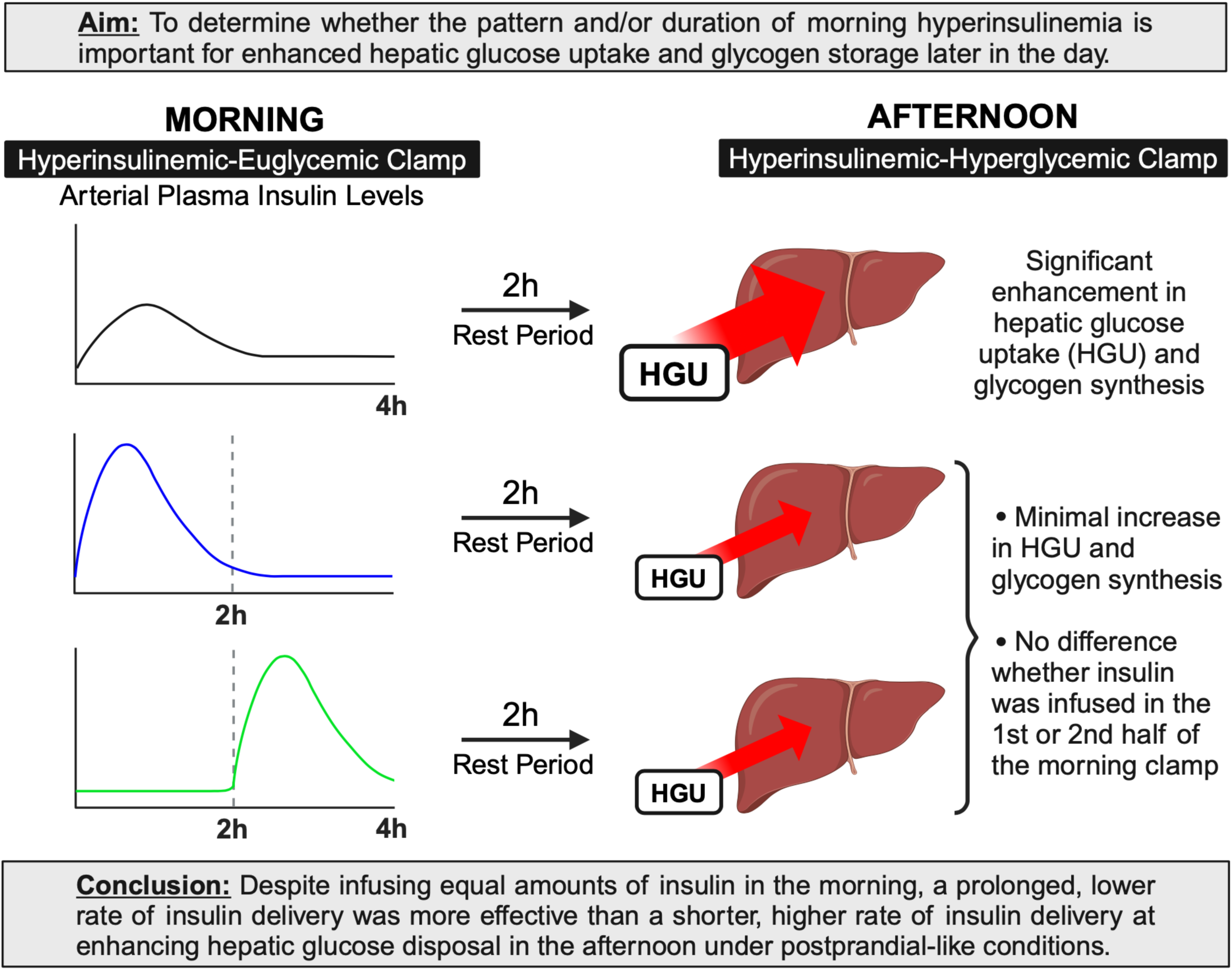
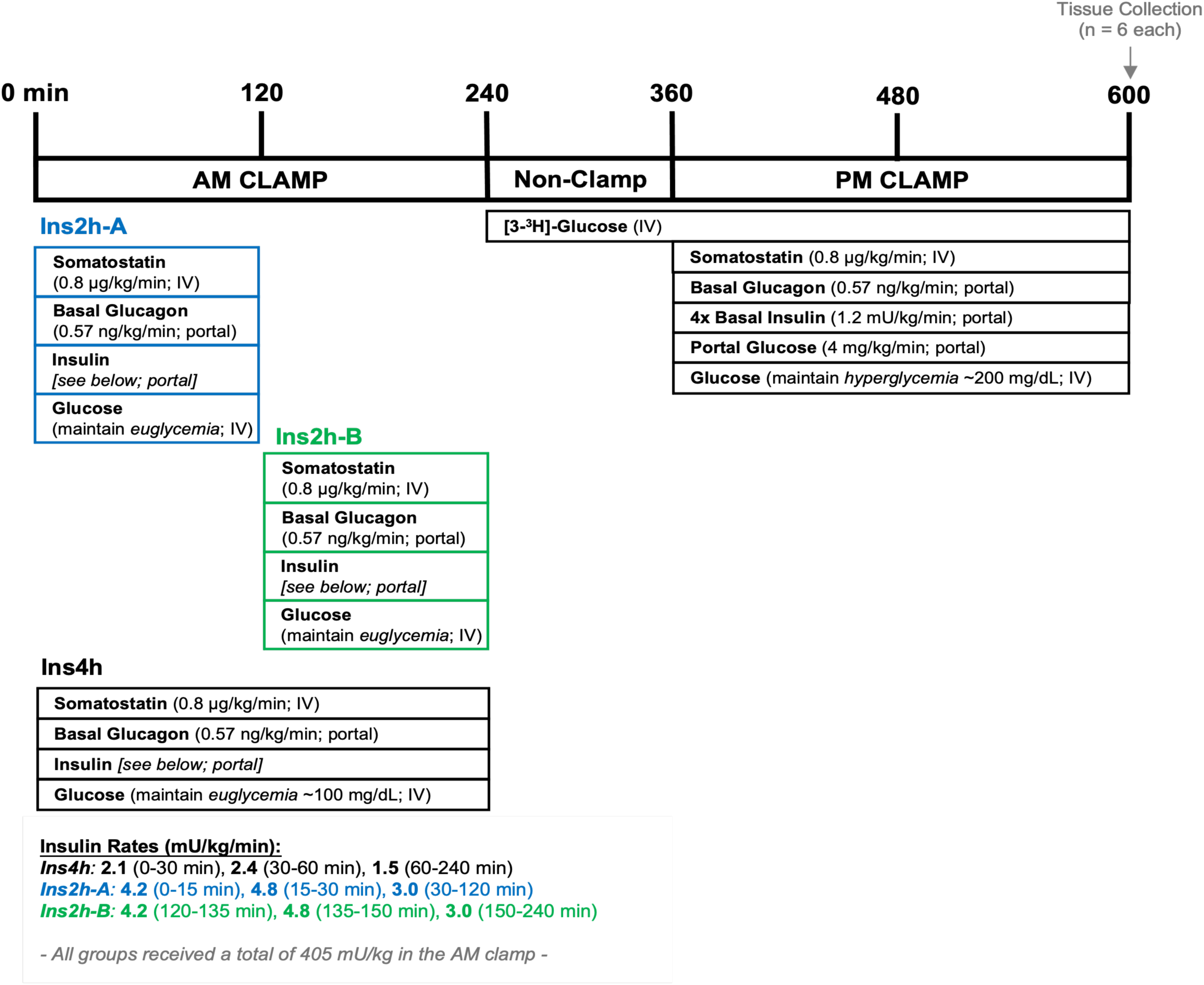
Study design. Dogs underwent two glucose clamp periods. All groups had a different pattern of insulin delivery for the morning (AM) hyperinsulinemic-euglycemic clamp, but ultimately received the same amount of insulin over a 2h or 4h period (405 mU/kg total insulin). Following the AM clamp, there was a 90-min tracer equilibration period (240 to 330 min) followed by 30 min of sampling under non-clamp conditions (330 to 360). Subsequently, all dogs underwent an afternoon (PM) hyperinsulinemic-hyperglycemic clamp with portal glucose delivery (360 to 600 min). Tissue collection occurred at the end of the PM clamp (600 min). Details can be found in the methods section. IV; intravenous infusion. Portal; portal vein infusion.

#### Non-clamp Period

In all groups, a primed, continuous infusion of [3-^3^H]-glucose (38 µCi prime and 0.38 μCi/min continuous rate; Revvitty, Waltham, MA) was administered via a peripheral leg vein beginning at 240 min. After a 90-minute tracer equilibration period, blood samples were collected from the arterial, portal, and hepatic sampling catheters at 330, 345, and 360 min to allow assessment of glucose kinetics before the PM clamp.

#### Afternoon (PM) Clamp

During the PM clamp, the liver was exposed to the three primary regulators that cause it to switch from fasting glucose production to mealtime glucose uptake and storage (i.e. hyperinsulinemia, hyperglycemia, and the negative arterial-to-portal vein glucose gradient that occurs when glucose is absorbed from the gut) (40, 41). All groups underwent the same PM hyperinsulinemic-hyperglycemic clamp from 360 to 600 min. Somatostatin was infused as described above, while glucagon (basal) and insulin (4x-basal; 1.2 mU/kg/min) were infused intraportally. 20% dextrose was infused into the hepatic portal circulation (4 mg/kg/min) and a primed, continuous infusion of 50% dextrose was administered intravenously via a leg vein to allow clamping of arterial blood glucose levels at ∼200 mg/dL. Thus, intraportal glucose and insulin infusions were used to mimic the postprandial conditions observed upon consumption of a moderate carbohydrate meal, but under steady state in a controlled environment.

Blood samples were collected throughout the experimental period from the artery, portal vein, and hepatic vein at 15-30 min intervals to allow measurement of hormones and substrates, which were analyzed as deemed appropriate. Arterial plasma glucose levels were monitored every 5 min during the AM clamp, non-clamp, and PM clamp periods. After obtaining the final blood sample, dogs were anesthetized while the hormone and glucose infusions were ongoing. Hepatic tissue was rapidly collected, flash-frozen in liquid nitrogen, and stored at -80°C.

### Analyses

#### Biochemical Methods

Arterial, hepatic portal vein, and hepatic vein whole blood samples collected during the study were analyzed using a variety of standard methods to obtain data regarding hormone and substrate balance across the liver. The arterial and portal blood samples were collected at the same time, followed by the hepatic venous blood sample. This allowed for the transit time of blood across the liver. Plasma glucose samples were immediately measured upon collection using a GM9 glucose analyzer (Analox Instruments Ltd., Stourbridge, UK). Blood lactate, glycerol, alanine, and non-esterified fatty acids were analyzed using enzymatic spectrophotometric methods **(see supplement).** Plasma insulin (#PI-12K, MilliporeSigma, Burlington, MA), glucagon (#GL-32K, MilliporeSigma), and cortisol (VUMC Analytical Services in-house primary antibody with I^125^ cortisol from MP Biomedicals, Santa Ana, CA) were measured in duplicate by radioimmunoassay. All samples were kept in an ice bath during the experiment and subsequently stored at - 80°C until assays were performed. For the determination of [3-^3^H]-glucose, plasma samples were deproteinized and quantified using liquid scintillation counting (42). Hepatic glycogen was quantitatively assessed using the Keppler and Decker amyloglucosidase method to determine total and radiolabeled glycogen **(see supplement).**

#### Molecular Analysis

Molecular methods were optimized to accurately assess enzyme activity of glucokinase (GK), glycogen synthase (GS), and glycogen phosphorylase (GP) through colorimetric and radioisotope techniques, total and phosphorylated protein levels for Akt (Ser473), GS, GP and total GK using Western blotting procedures, and mRNA of GK, G6Pase, and PEPCK by RT-PCR **(see supplement).** Basal liver samples from 3 overnight-fasted dogs maintained on a standard chow diet that had undergone no experimental intervention were utilized for reference purposes.

#### Calculations

Direct net hepatic glucose balance was calculated using the A-V difference method using the equation

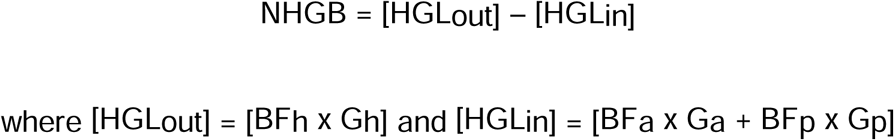

HGLout represents the glucose load exiting the liver, and HGLin is the glucose load entering the liver. BF represents blood flow, G represents the blood glucose concentration, and A, P, and H represent the hepatic artery, hepatic portal vein, and hepatic vein. Under conditions of hepatic glucose production, NHGB is positive, whereas it is negative when the liver is in uptake mode. Direct net hepatic balance of other substrates across the liver may be calculated using this method as well. To avoid any errors in calculating NHGB due to possible incomplete mixing of substrates infused into the hepatic portal vein, HGLin was also calculated using an indirect method:

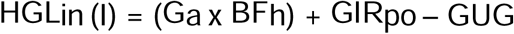

Where GIR_PO_ is the intraportal glucose infusion rate and GUG is the uptake of glucose by the gastrointestinal tract. GUG was calculated as

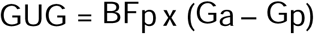

Thus, indirect NHGB = HGLout– HGLin (I). There were no significant differences between direct and indirect net hepatic balance for any of the substrates measured. The data in this report were calculated using the direct method. Hepatic sinusoidal hormones and substrates were determined using the equation for HGLin. The fractional extraction of a hormone or substrate by the liver was calculated as the net hepatic balance ÷ HLin. Absolute (unidirectional) HGU was determined by multiplying the fractional extraction of [3-^3^H]-glucose (which is not subject to portal vein mixing errors since the tracer was infused intravenously into a leg vein) by HGLin. Non-hepatic glucose uptake (non-HGU) was determined by subtracting HGU and the change in glucose mass in the blood from the total GIR.

Plasma glucose concentrations were converted to blood concentrations using a conversion factor established previously (43). Net hepatic carbon retention (NHCR), a proxy for glycogen synthesis over time, was calculated as

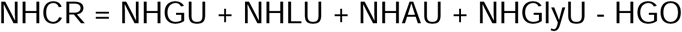

Where NHGU represents net hepatic glucose uptake, NHLU represents net hepatic uptake of lactate, NHAU represents net hepatic uptake of alanine, NHGlyU represents net hepatic uptake of glycerol, and HGO represents hepatic glucose oxidation, which was estimated to be 0.2 mg/kg/min throughout the experiment as determined in earlier studies (44). Lactate, alanine, and glycerol were converted to glucose equivalents before calculating NHCR. NHAU was doubled to account for net hepatic uptake of all amino acids, as alanine accounts for approximately 50% of gluconeogenic amino acid precursor uptake (45, 46). Direct glycogen synthesis was calculated by taking the total amount of radiolabeled glycogen (tracer counts liberated using Keppler and Decker method, **see supplement**) and dividing it by the specific activity (SA) inflow of the precursor pool which was calculated as

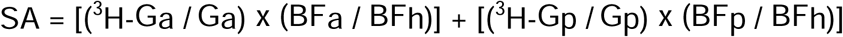

Where ^3^H-G represents tracer-determined blood ^3^H glucose.

#### Statistical Analysis

Data are expressed as mean ± SEM. Statistical comparisons within and between groups were carried out using a two-way analysis of variance with repeated measures design. One-way analysis of variance was used for temporal assessment within a single group. Post hoc analysis was performed using the Student-Newman-Keuls multiple comparisons test. All statistics were analyzed using GraphPad Prism software. Respective areas under the curve (AUCs) were compared using one-way analysis of variance with Tukey’s post hoc analysis. A P value <0.05 was considered statistically significant.

## Results

### AM Clamp Data

Plasma glucose concentrations did not differ between any of the groups before the start of the AM clamp, and euglycemia was maintained in all dogs during the AM clamp **(Fig. 2A).** During the first 120 min, arterial insulin levels were significantly greater in the Ins2h-A group in comparison to the Ins4h group **(Fig. 2B).** At 120 min, insulin infusion ceased in the Ins2h-A group and began in the Ins2h-B group. From 120 to 240 min, Ins2h-B achieved arterial insulin levels similar to those in Ins2h-A. The calculated areas under the curve for Ins2h-A and Ins2h-B were 2-fold greater than that of Ins4h, with no significant difference between the 2h groups **(Fig. 2C).** The glucose infusion rate (GIR) for each group reached comparable maximum rates, however, the GIR for Ins4h was sustained throughout the entire AM clamp, then decreased, whereas GIR decreased after the end of insulin infusion at 120 min in Ins2h-A and after 240 min in Ins2h-B **(Fig. 2D).** The ΔAUC for GIR was 2-times greater in Ins4h than Ins2h-A or Ins2h, despite the ΔAUC for insulin being half as great in Ins4h **(Fig. 2E)**. Glucose infusion continued in Ins4h and Ins2h-B after the end of the AM clamp (during the non-clamp period) to maintain euglycemia until exogenous glucose was no longer needed (terminated in all groups by 330 min).

**Figure 2.**
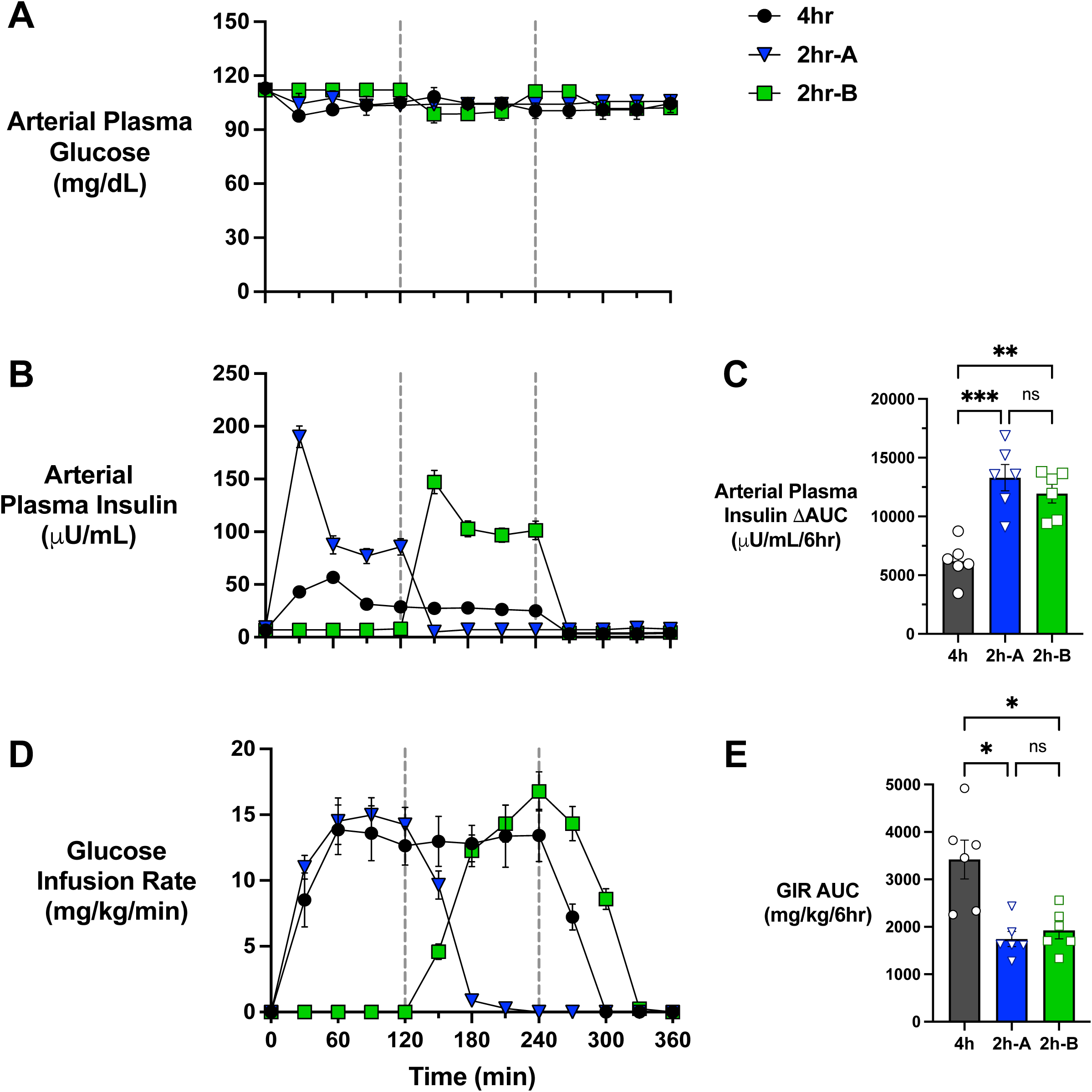
AM hyperinsulinemic-euglycemic clamp data. A vertical line at 120 min marks the end of the Ins2h-A and the start of the Ins2h-B clamp period. Arterial plasma glucose (A) and insulin (B) concentrations, as well as glucose infusion rates (D) required to maintain euglycemia over the course of the 4h morning period, are shown for Ins4h, Ins2h-A, and Ins2h-B; *n*=6/group. Bar graphs indicate the area under the curve (AUC) for arterial insulin concentrations (C) and glucose infusion rates (E) during the 4h morning period in each dog. Data are expressed as mean ± SEM. *P<0.05, **P<0.01 between groups. All comparisons that did not reach statistical significance are denoted ns.

### PM Clamp Data

During the PM clamp, the liver was exposed to the three primary mealtime regulators of HGU. In all groups, the plasma glucose levels were doubled **(Fig. 3A, 3B, 3D)**, a negative arterial-to-portal vein glucose gradient was established **(Fig. 3C),** and insulin increased 4-fold **(Fig. 3E, 3F)**, as intended. Additionally, the arterial and hepatic sinusoidal plasma glucagon concentrations were kept at basal throughout the PM clamp in all groups. There were no differences in arterial cortisol levels between groups, indicating the animals were not stressed, and these levels remained low throughout the experiment **(Table 1)** (47, 48).

**Figure 3.**
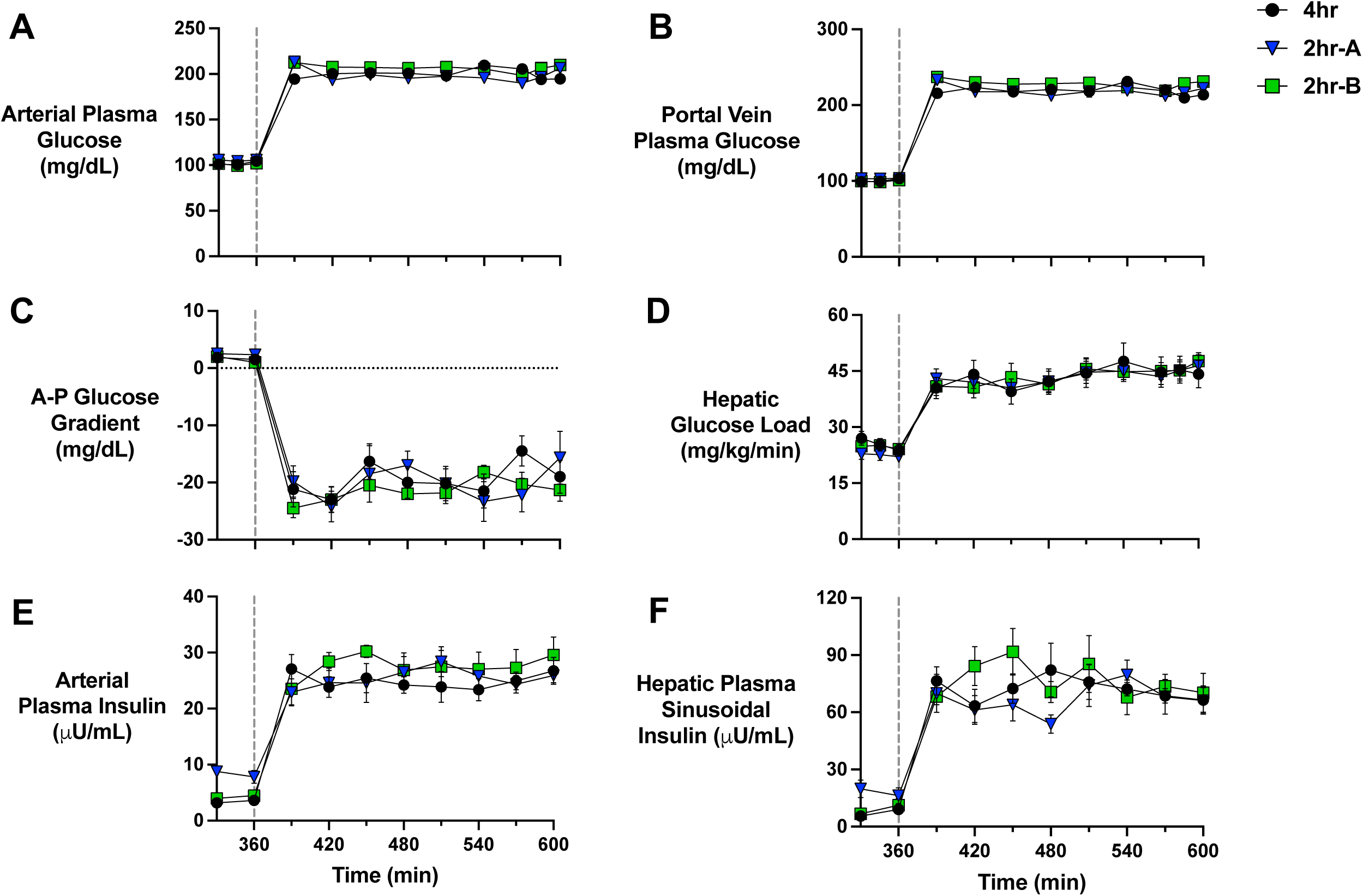
PM clamp glucose and hormone data. A vertical line at 360 min marks the start of the afternoon (PM) clamp. Arterial plasma glucose (A), portal vein plasma glucose (B), the difference between the artery and the portal vein plasma glucose levels (C), hepatic glucose load (D), arterial plasma insulin (E), and plasma insulin at the hepatic sinusoids (F) during the last 30 min of the non-clamp and 4h PM clamp period are shown for Ins4h, Ins2h-A, and Ins2h-B; *n*=6/group. All groups were matched during the PM clamp for these parameters. Data are expressed as mean ± SEM.

**Table 1:**
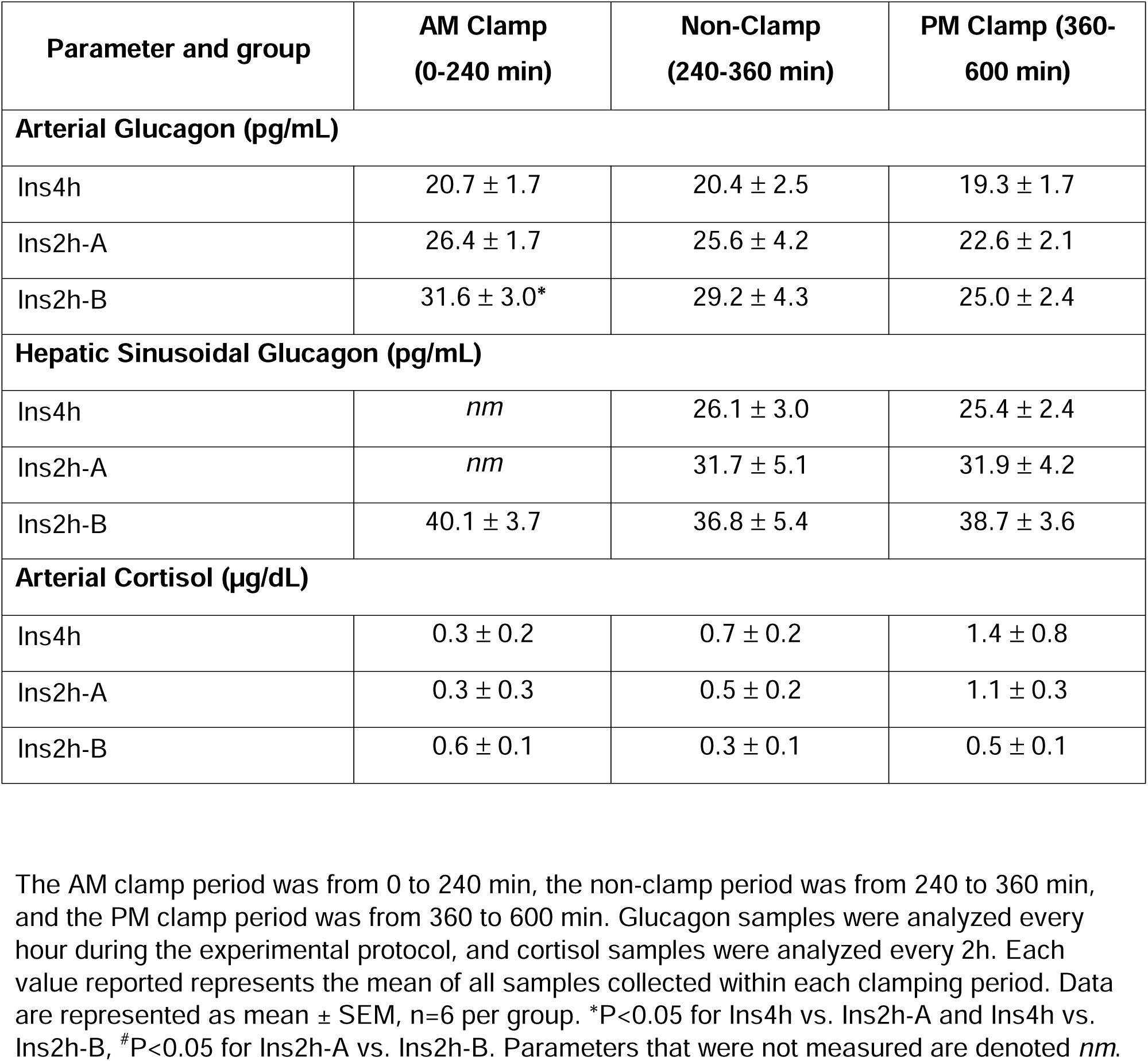
Average plasma hormone concentrations during the AM clamp, non-clamp, and PM clamp periods.

The total average GIR (portal and leg vein glucose infusion) over the PM clamp trended higher in Ins4h vs. Ins2h-A and Ins2h-B (17.7 ± 2.3 vs. 13.7 ± 2.4 and 13.3 ± 1.2 mg/kg/min, respectively), but this did not reach significance **(Fig. 4A).** The ΔAUC for GIR also tended to be greater in Ins4h vs. Ins2h-A and Ins2h-B (3621 ± 499 vs. 2743 ± 453 and 2655 ± 253 mg/kg/4hr, respectively; **Fig. 4B**). All dogs rapidly switched to a state of HGU at the onset of the PM clamp. HGU was 40% and 30% greater in Ins4h compared to Ins2h-A and Ins2h-B, respectively (ΔAUC for HGU 1393 ± 199 vs. 822 ± 68 and 900 ± 76 mg/kg/4hr, respectively, P<0.05) with no difference between the 2h groups **(Fig. 4C, 4D).** Non-HGU, which under these conditions is primarily attributed to skeletal muscle glucose uptake, was indistinguishable between groups throughout the PM clamp, although there was a tendency for non-HGU to be slightly greater in the Ins4h group **(Fig. 4E, 4F).** In all groups, muscle glucose uptake accounted for 55-58% of total glucose disposal during the 4h PM clamp.

**Figure 4.**
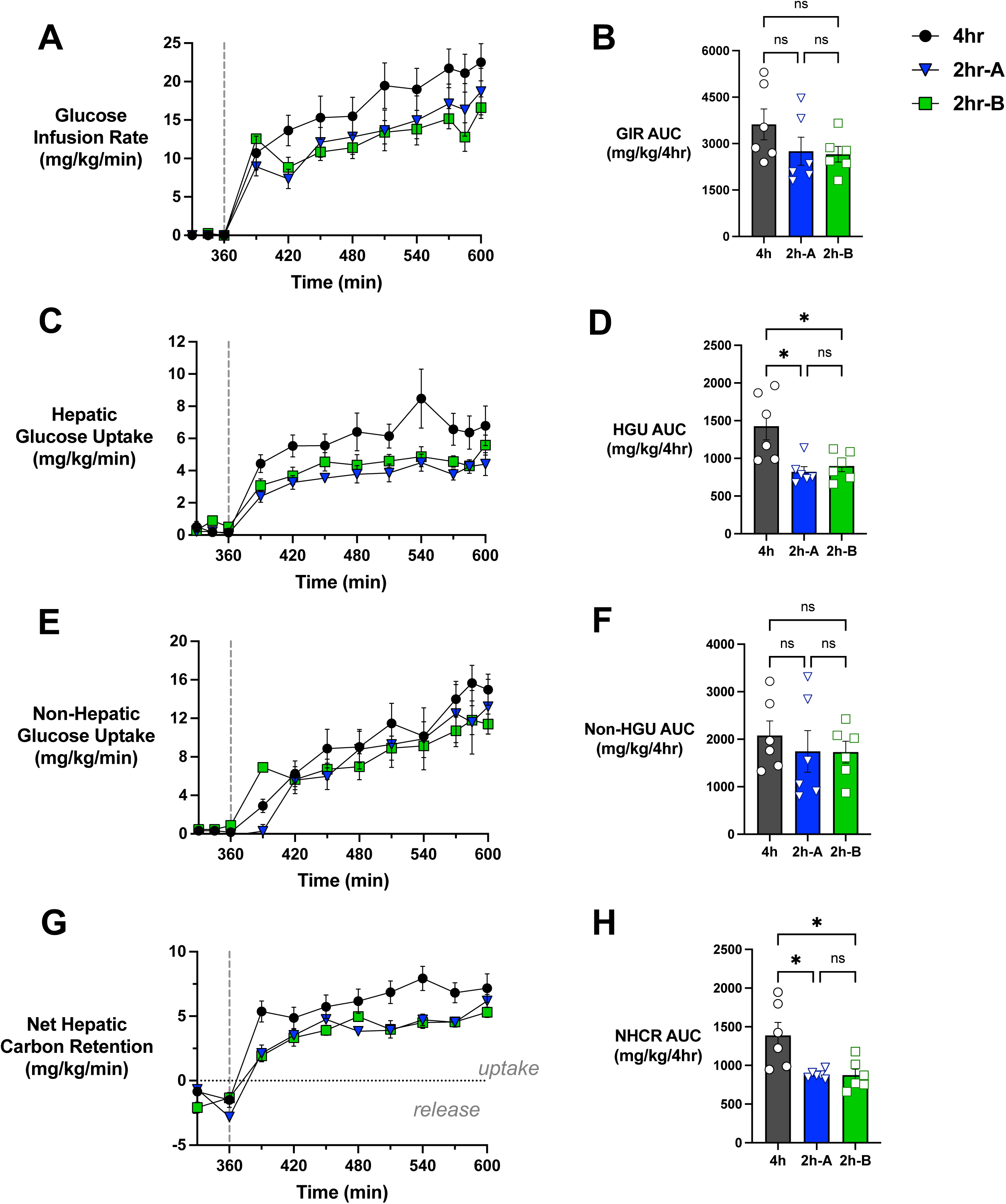
PM hyperinsulinemic-hyperglycemic clamp glucose uptake. A vertical line at 360 min marks the start of the afternoon (PM) clamp. Glucose infusion rate (A), hepatic glucose uptake (C), non-hepatic glucose uptake (E), and net hepatic carbon retention (G) are shown over time. Their respective areas under the curve (B, D, F, H) are shown for the non-clamp and PM clamp period for Ins4h, Ins2h-A, and Ins2h-B; *n*=6/group. *P<0.05, **P<0.01 between groups. ns = not significant.

Net hepatic carbon retention (NHCR) was significantly greater in Ins4h than in Ins2h-A or Ins2h-B (**Fig. 4G,** P<0.013). This rate includes glucose and all 3-carbon precursors taken up by the liver, including glycerol, amino acids, and lactate. Previous studies done in our lab have shown that de novo hepatic lipogenesis is a relatively small component of hepatic glucose disposal under similar conditions (37, 49). The NHCR AUC for Ins2h-A and Ins2h-B were both 63% of the AUC for Ins4h **(Fig. 4H).** At the onset of the PM clamp, net hepatic non-esterified fatty acid, glycerol, and alanine uptake decreased and remained suppressed **(Table 2).** All dogs switched to net hepatic lactate output and remained in that state for the entirety of the PM clamp, with no differences between groups **(Table 2).**

**Table 2:** PM clamp non-esterified fatty acid and metabolite flux data.

| Parameter and group | Non-Clamp | PM Clamp |  |  |  |  |
| --- | --- | --- | --- | --- | --- | --- |
|  | 360 | 420 | 480 | 540 | 600 | PM Clamp Mean |
| **Arterial Plasma NEFA (μmol/L)** |  |  |  |  |  |  |
| Ins4h | 828±109 | 113±37 | 77±30 | 67±22 | 46±6 | 80±17 |
| Ins2h-A | 960±108 | 165±44 | 83±28 | 71±22 | 56±19 | 128±40 |
| Ins2h-B | 1127±140 | 102±23 | 86±18 | 50±3 | 86±15* | 100±17 |
| **Net Hepatic NEFA Uptake (μmol/kg/min)** |  |  |  |  |  |  |
| Ins4h | 2.1±0.3 | 0.4±0.2 | 0.4±0.2 | 0.2±0.1 | 0.3±0.1 | 0.3±0.1 |
| Ins2h-A | 1.9±0.2 | 0.5±0.1# | 0.2±0.1 | 0.3±0.2 | 0.4±0.1 | 0.4±0.1 |
| Ins2h-B | 2.1±0.4 | 0.3±0.1 | 0.2±0.1 | 0.1±0.1 | 0.2±0.1 | 0.2±0.1 |
| **Arterial Blood Glycerol (μmol/L)** |  |  |  |  |  |  |
| Ins4h | 88±12 | 26±5 | 22±2 | 19±4 | 17±4 | 23±3 |
| Ins2h-A | 92±6 | 37±5 | 33±6 | 27±5 | 26±5 | 32±5 |
| Ins2h-B | 123±15 | 28±5 | 25±5 | 29±7 | 21±4 | 27±5 |
| **Net Hepatic Glycerol Uptake (μmol/kg/min)** |  |  |  |  |  |  |
| Ins4h | 2.0±0.4 | 0.4±0.1 | 0.3±0.1 | 0.2±0.1* | 0.2±0.1 | 0.3±0.1 |
| Ins2h-A | 2.0±0.2# | 0.8±0.1* | 0.6±0.1 | 0.6±0.1 | 0.6±0.1* | 0.6±0.1 |
| Ins2h-B | 2.8±0.3 | 0.4±0.1 | 0.4±0.1 | 0.6±0.1 | 0.3±0.1# | 0.5±0.1 |
| **Arterial Blood Lactate (μmol/L)** |  |  |  |  |  |  |
| Ins4h | 659±169 | 781±55 | 656±85 | 770±103 | 841±92 | 813±77 |
| Ins2h-A | 323±25 | 848±94 | 673±84 | 798±82 | 824±75 | 776±74 |
| Ins2h-B | 380±40 | 796±82 | 658±72 | 710±50 | 753±68 | 711±45 |
| **Net Hepatic Lactate Output (μmol/kg/min)** |  |  |  |  |  |  |
| Ins4h | -2.8±2.8 | 16.0±1.8 | 10.4±2.4 | 11.6±2.6 | 10.7±2.4 | **12.3±2.1** |
| Ins2h-A | -3.2±0.8 | 14.2±3.3 | 7.9±2.3 | 7.0±2.0 | 6.7±1.9 | **9.8±2.4** |
| Ins2h-B | -1.1±1.8 | 15.8±2.5 | 9.9±1.7 | 9.7±1.7 | 8.8±2.4 | **11.3±1.8** |
| **Arterial Blood Alanine (μmol/L)** |  |  |  |  |  |  |
| Ins4h | 265±39 | 282±25 | 246±19 | 234±15 | 220±17 | **250±20** |
| Ins2h-A | 224±15 | 276±18 | 244±27 | 233±30 | 221±29 | **245±27** |
| Ins2h-B | 223±10 | 243±15 | 238±19 | 278±55 | 217±15 | **238±14** |
| **Net Hepatic Alanine Uptake (μmol/kg/min)** |  |  |  |  |  |  |
| Ins4h | 2.4±0.5 | 2.1±0.2 | 1.8±0.1 | 1.8±0.2 | 1.7±0.3 | **2.0±0.2** |
| Ins2h-A | 1.8±0.2 | 1.3±0.2* | 1.7±0.2 | 1.6±0.1 | 1.5±0.2 | **1.4±0.1** |
| Ins2h-B | 1.8±0.3 | 1.5±0.2 | 1.4±0.1 | 2.2±0.5 | 1.1±0.4 | **1.5±0.1** |

The non-clamp period was from 240 to 360 min, and the PM clamp period was from 360 to 600 min. Average group values are presented at the onset of the clamp (360 min) and every hour during the PM clamp. Blood samples were collected every 30 min during the non-clamp and PM clamp periods. Data are represented as mean ± SEM, n=6 per group. \*P<0.05 for Ins4h vs. Ins2h-A and Ins4h vs. Ins2h-B, #P<0.05 for Ins2h-A vs. Ins2h-B.

### Liver tissue analyses

Compared to basal overnight fasted dogs, Akt phosphorylation was elevated similarly in all groups at the end of the PM clamp **(Fig. 5A).** Glucokinase (GK) mRNA levels were not significantly different between the three groups, although GK transcript levels were greatest in the Ins4h group and lowest in the Ins2h-B group **(Fig. 5B).** Furthermore, GK protein expression and activity were elevated in all three groups compared to basal in a trend similar to changes in GK mRNA **(Fig. 5C, 5D).** These data suggest that the increase in PM HGU in the Ins4h group compared to the 2h groups does not appear to be due to an acute increase in insulin receptor activation or GK transcription, protein levels, or enzyme activity.

**Figure 5.**
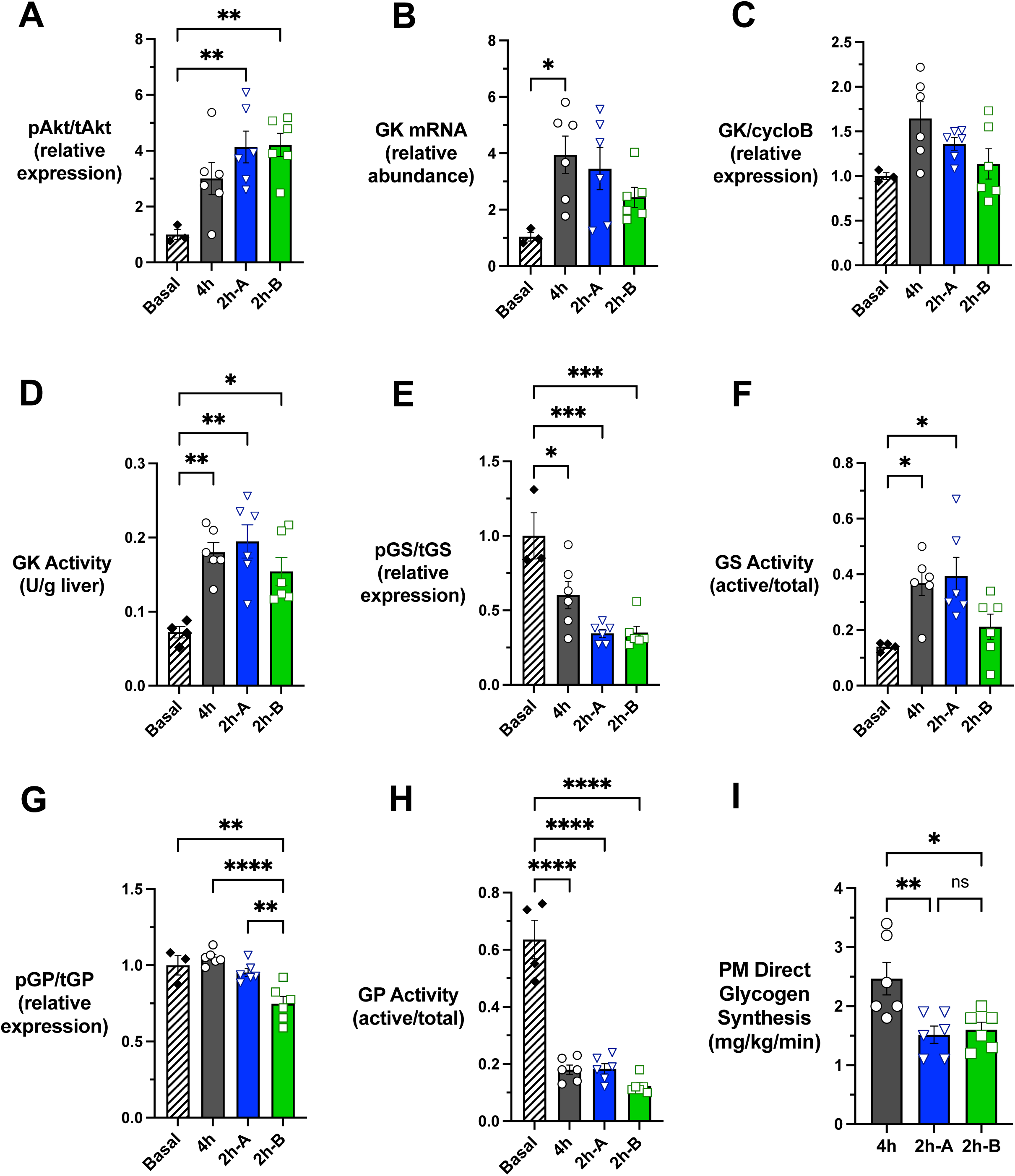
Liver tissue analyses. Phosphorylated Akt protein (A), glucokinase (GK) mRNA (B), GK protein (C), GK activity (D), phosphorylated glycogen synthase (GS) protein (E), GS activity (F), phosphorylated glycogen phosphorylase (GP) protein (G), GP activity (H), and the rate of direct glycogen synthesis during the afternoon (PM) clamp (I) are shown for basal (when applicable, *n=3-4)*, as well as Ins4h, Ins2h-A, and Ins2h-B (*n*=6/group). Data are expressed as mean ± SEM. *P<0.05, **P<0.01, ***P<0.001, ****P<0.0001 between groups. All other between-group comparisons not denoted with a P-value are not significant (ns).

When assessing the enzymes involved in glycogen metabolism, all three groups had greater dephosphorylation and thus activation of glycogen synthase (GS) when compared to baseline **(Fig. 5E)**. The greatest GS activity was observed in the Ins4h and Ins2h-A groups when compared to basal, although there was a strong tendency for Ins2h-B to be lower than Ins4h and Ins2h-A (P<0.074, **Fig. 5F)**. Additionally, the Ins2h-B group had significantly greater dephosphorylation and thus deactivation of glycogen phosphorylase (GP) when compared to the other groups **(Fig. 5G).** There were no apparent differences between any of the groups regarding the suppression of GP activity **(Fig. 5H).** The rate of direct glycogen synthesis during the PM clamp, which is the amount of radiolabeled glucose being shuttled into glycogen, was 1.6-fold greater in Ins4h vs. Ins2h-A and Ins2h-B (mean direct glycogen synthesis of 2.5 ± 0.3 mg/kg/min in Ins4h vs. 1.5 ± 0.1 mg/kg/min in Ins2h-A and 1.6 ± 0.1 mg/kg/min in Ins2h-B, **Fig. 5I).** There was a tendency for the molecular responses in the Ins2h-B group to be somewhat less receptive to the postprandial signals during the PM clamp compared to the Ins4h and Ins2h-A groups, although not statistically significant (except for GP phosphorylation).

## Discussion

Although the second meal phenomenon was identified more than a century ago, the factors causing it have remained poorly understood (50). It has been hypothesized that the lower glycemic response observed upon consuming a second glucose load could be due to an enhancement in muscle insulin sensitivity, enhanced insulin secretion, a reduction in the absorption of oral glucose mediated by increased GLP-1 secretion, or a combination of these factors (27, 29, 51). Our current and previous data, along with that of others, do not support the involvement of any of these factors in the phenomenon (27, 32, 37, 52). Rather, our recent studies determined that the second meal phenomenon was primarily due to enhanced HGU secondary to elevated insulin exposure in the morning (32). This raises the question of how morning insulin delivery might be tailored to maximize this response. We sought to address this issue by determining the importance of 1) the duration of AM insulin exposure, 2) the magnitude of AM insulin exposure, and 3) the timing between AM insulin exposure and the PM clamp (mimicking a second meal).

In the current set of experiments, we observed that 4h of morning hyperinsulinemia was more effective than 2h of morning hyperinsulinemia at priming the liver for enhanced PM hepatic glucose uptake and hepatic glycogen synthesis during a subsequent hyperinsulinemic-hyperglycemic clamp. The rates of AM insulin delivery were designed to match the total amount of insulin delivered between all three groups yet permit a shorter duration and increased magnitude of insulin delivery in the Ins2h groups (Ins4h AM insulin delivery rates were doubled and delivered over half the amount of time in the Ins2h groups). It is evident that a prolonged, lower rate of AM insulin delivery improved PM liver glucose metabolism more effectively than a shorter, higher rate of AM insulin delivery. These results are consistent with findings in healthy adults indicating that slow-release carbohydrates, compared with more rapidly absorbed foods consumed at breakfast, improved the response to a standardized lunch (53). In that investigation, the glycemic pattern was mimicked by carefully matched feeding over a 4h period, but insulin responses were not matched throughout the morning period. Thus, those findings do not provide information about the impact of the pattern of the insulin response throughout the morning period, which we show in this report.

In a former set of experiments, we mimicked “breakfast-skipping” by infusing saline over 4h in the morning (37). We found that during a subsequent 4h PM HIHG clamp identical to the PM clamp in this study, PM HGU was 2.9 ± 0.2 mg/kg/min in this “breakfast-skipping” group (37). Although PM HGU was lower in the 2h groups in this study compared to the Ins4h group (3.7 ± 0.3 and 4.4 ± 0.4 vs. 6.3 ± 0.9 mg/kg/min in Ins2h-A and Ins2h-B vs. Ins4h, respectively), it was still enhanced relative to the “breakfast-skipping” group, which serves as a negative second meal control. Thus, even though the 2h duration of insulin exposure in the morning did not achieve the rate of PM HGU observed in the 4h group, it still had a stimulatory effect. Remarkably, the 2h groups were incredibly similar when assessing the PM flux data, regardless of whether there was 2h or 4h of insulin withdrawal (no significant differences between the 2h groups for any flux parameter measured). Given that there was no sign of waning when the rest period was extended from 2h to 4h, it remains to be determined how long the insulin priming effect persists.

We next sought to understand the molecular mediators of these effects. When assessing insulin action, a determinant of the PM HGU response, the duration of AM hyperinsulinemia did not appear to significantly affect the insulin receptor or downstream signaling in the afternoon. Akt phosphorylation was elevated similarly in all groups, at least at the end of the study, suggesting that the enhancement in PM HGU in the 4h group compared to the 2h groups was not due to increased PM insulin action. Despite differences in hepatic glycogen content, the activation of signals mediating glycogen storage does not appear to be significantly different between the 4h or 2h groups. Glycogen synthase (GS) and glycogen phosphorylase (GP) activity and protein levels are rate-limiting steps in hepatic glycogen synthesis and breakdown (54, 55), and GS and GP activities were not significantly different between groups at the time of tissue collection. Additionally, there were no significant differences in PM GK mRNA, GK protein, or GK activity, although all were elevated compared to baseline in all three experimental groups. It should be noted that since tissues were collected at the end of the PM period, differential effects of morning insulin priming on molecular mediators of the effect at the start of the PM period could have been obscured by the subsequent hyperinsulinemic-hyperglycemic clamp. One possibility is that there was a difference in GK translocation between the 4h and 2h groups that is undetectable using current methods. In the fasted state, GK is complexed with glucokinase regulatory protein (GKRP) and sequestered in the nucleus in its inactive state. However, in postprandial conditions (such as the elevated glucose conditions in the PM HIHG clamp), GK quickly dissociates from GKRP and moves to the cytoplasm in its active form (56, 57). GKRP functions both as a metabolic sensor and as a nuclear chaperone and is, therefore, detected in both nuclear and cytosolic fractions (58). It has been shown previously that GK rapidly translocates in a manner that correlates with changes in HGU, supporting the hypothesis that GK translocation could be increased in the 4h group compared to the 2h groups (59). In the past, we have employed immunohistochemical staining and confocal microscopy procedures in rodent experiments to track GK translocation over time using fresh hepatic tissue under a variety of experimental conditions (59). Our lab has previously attempted to quantify GK translocation using similar methods in the dog, but has remained unsuccessful in doing so, largely due to the lack of canine-specific antibodies.

Consequent to the presence and activation of GK, elevated levels of glucose-6-phosphate elicit a synergistic effect, along with glucose, in promoting GS activation and GP inactivation through allosteric control (60, 61). Furthermore, glucose-6-phosphate and glucose have been suggested to impact the cellular localization and activity of these enzymes, which could, in part, potentially explain the increase in PM hepatic glycogen storage in the Ins4h group (60, 62, 63). It is noteworthy that our GS and GP activity assays reflect the phosphorylation state of these enzymes, and do not necessarily take into account the allosteric activation of GS or inactivation of GP that do not involve a phosphorylation event (61). These allosteric effects may further explain the increase in PM HGU and NHCR in Ins4h compared to the 2h groups which is not further supported or explained by observed differences in GS and GP activities.

While there do not appear to be significant molecular changes between the 2h groups aside from GP phosphorylation, it is important to note that shortening the priming washout period from 4 to 2h seems to have had a subtle effect on GK transcription, protein, and activity, which all trended lower ins Ins2h-B than observed in the Ins4h and Ins2h-A groups. Furthermore, although GS activity was lower in Ins2h-B, GP activity was as well. The ratio of GS to GP activity suggests the effects on these enzymes somewhat cancel each other out, which could explain why there did not appear to be an impact on PM hepatic glycogen synthesis in Ins2h-B relative to Ins2h-A. The variance observed in the molecular data makes it difficult to draw any solid conclusions about how the timing between AM insulin exposure and the PM HIHG clamp might be impacting the transcription and function of key enzymes involved in glucose uptake and glycogen metabolism during the PM.

The total amount of insulin delivered in the morning was identical in the Ins4h and both 2h groups, yet the insulin AUCs were twice as great in the 2h groups. The peak insulin levels in both 2h groups (30 min after the start of the AM clamp) were ∼4-fold greater than the peak insulin level in the 4h group, despite both 2h groups only having a doubling in the insulin infusion rates compared to the 4h group over the 2h infusion period. The capacity for insulin clearance by the kidneys and liver, the main organs responsible for removing insulin from the bloodstream, can become saturated, which likely explains the observed differences in insulin levels (64–66). However, it is important to recognize that despite the greater rate of AM insulin delivery in both 2h groups, the glucose infusion rates required to maintain euglycemia were the same as those seen in the 4h group, thus the lower levels of insulin (as seen in the 4h group) were sufficient to maximally stimulate glucose uptake in the morning. This suggests that the rate of insulin delivery in the AM clamp in the 2h groups was less than optimal for priming the liver for enhanced PM HGU. It will be important in future studies to assess how a lower dose of morning insulin might impact the effects observed in the PM HIHG clamp.

In conclusion, the duration of morning insulin delivery is a critical regulator of HGU and glycogen synthesis during a second glucose load. When the same amount of insulin was administered in the morning, 4h of lesser hyperinsulinemia had a greater effect on HGU and hepatic glycogen synthesis in the afternoon than did 2h of greater hyperinsulinemia, independent of whether the amount of time elapsed between AM and PM periods was 2 or 4h. It is evident that the liver is highly responsive to changes in the rate and pattern of insulin delivery, with the levels attained in the Ins4h group prompting a robust effect on hepatic glucose uptake and storage in the afternoon. The results of this study provide valuable insight into how meal composition and size might be adapted to improve glycemic control and maximize hepatic glycogen stores. Considering that glucose homeostasis is largely dysregulated in individuals with insulin-dependent diabetes, the results from this study give insight into how a morning meal might be able to improve blood sugar regulation throughout the remainder of the day. Delivering a lower rate of insulin in the morning over an extended period versus a greater rate of insulin over a shorter period should prompt greater HGU and glycogen storage at lunchtime, and perhaps even later meals. Additionally, our data provide metabolic support for the consumption of slower absorbing carbohydrates during meals.

## Data Availability

Supplemental material is publically available via the following DOI for Figshare: doi.org/10.6084/m9.figshare.26791330.v1. The data generated during and analyzed during the current study are available from the corresponding author upon reasonable request.

## Grants

This work was supported by National Institutes of Health (NIH) Grant R01DK131082. H.W. was supported in part by the Vanderbilt University Training Program in Molecular Endocrinology NIH grant 5T32DK007563. Hormone analysis was completed by Vanderbilt’s Hormone Assay and Analytical Services Core, supported by NIH grants DK020593 and DK059637. Vanderbilt’s Large Animal Core provided surgical expertise, supported by the NIH Grant DK020593.

## Disclosures

A.D.C. has research contracts with Fractyl, Abvance, Fluidics, and Novo Nordisk, as well as grants from the NIH and the Helmsley Charitable Trust. A.D.C. is a consultant to Novo Nordisk, Paratus, Portal Insulin, Fractyl, and Thetis. No other individuals have conflicts of interest relevant to this article.

## Author Contributions

H.W., M.C.M., and A.D.C. participated in the design of experiments; H.W. and M.C.M. directed all experiments, collected, and interpreted data; M.S.S, B.F., M.S., and D.S.E. participated in the experiments; B.F. was responsible for surgical preparation and oversight of animal care; M.S.S. carried out biochemical and tissue analysis; H.W., D.S.E., and A.D.C. interpreted experimental results; H.W. prepared figures and drafted the manuscript; H.W., D.S.E., and A.D.C. edited and revised the manuscript; All authors approved the final version of the manuscript. A.D.C. is the guarantor of this work and, as such, has full access to all the data in the study and takes responsibility for the integrity of the data and the accuracy of the data analysis.

## Acknowledgements

Graphical abstract was created with Bio Render and published with permission.

## Notes

https://doi.org/10.6084/m9.figshare.26791330.v1

### Competing Interest Statement

Conflict of Interest Statement: Hannah Waterman, Mary Moore, Marta Smith, Ben Farmer, Melanie Scott, and Dale Edgerton have no conflicts of interest. Alan D. Cherrington has the following potential conflicts of interest to disclose: A.D.C. has research contracts with Fractyl, Abvance, Fluidics, and Novo Nordisk, as well as grants from the NIH and the Helmsley Charitable Trust. A.D.C. is a consultant to Novo Nordisk, Paratus, Portal Insulin, Fractyl, and Thetis.

### Summary of Updates

The manuscript was updated to reflect changes during the peer-review process.

## References

1. Li ZH, Xu L, Dai R, Li LJ, and Wang HJ. Effects of regular breakfast habits on metabolic and cardiovascular diseases: A protocol for systematic review and meta-analysis. Medicine (Baltimore) 100: e27629, 2021.

2. Ma X, Chen Q, Pu Y, Guo M, Jiang Z, Huang W, Long Y, and Xu Y. Skipping breakfast is associated with overweight and obesity: A systematic review and meta-analysis. Obes Res Clin Pract 14: 1–8, 2020.

3. Ballon A, Neuenschwander M, and Schlesinger S. Breakfast Skipping Is Associated with Increased Risk of Type 2 Diabetes among Adults: A Systematic Review and Meta-Analysis of Prospective Cohort Studies. J Nutr 149: 106–113, 2019.

4. Chen H, Zhang B, Ge Y, Shi H, Song S, Xue W, Li J, Fu K, Chen X, Teng W, and Tian L. Association between skipping breakfast and risk of cardiovascular disease and all cause mortality: A meta-analysis. Clin Nutr 39: 2982–2988, 2020.

5. Ogata H, Hatamoto Y, Goto Y, Tajiri E, Yoshimura E, Kiyono K, Uehara Y, Kawanaka K, Omi N, and Tanaka H. Association between breakfast skipping and postprandial hyperglycaemia after lunch in healthy young individuals. Br J Nutr 122: 431–440, 2019.

6. Rong S, Snetselaar LG, Xu G, Sun Y, Liu B, Wallace RB, and Bao W. Association of Skipping Breakfast With Cardiovascular and All-Cause Mortality. Journal of the American College of Cardiology 73: 2025–2032, 2019.

7. St-Onge MP, Ard J, Baskin ML, Chiuve SE, Johnson HM, Kris-Etherton P, and Varady K. Meal Timing and Frequency: Implications for Cardiovascular Disease Prevention: A Scientific Statement From the American Heart Association. Circulation 135: e96–e121, 2017.

8. Ferrer-Cascales R, Sánchez-SanSegundo M, Ruiz-Robledillo N, Albaladejo-Blázquez N, Laguna-Pérez A, and Zaragoza-Martí A. Eat or Skip Breakfast? The Important Role of Breakfast Quality for Health-Related Quality of Life, Stress and Depression in Spanish Adolescents. Int J Environ Res Public Health 15: 2018.

9. Kahleova H, Petersen KF, Shulman GI, Alwarith J, Rembert E, Tura A, Hill M, Holubkov R, and Barnard ND. Effect of a Low-Fat Vegan Diet on Body Weight, Insulin Sensitivity, Postprandial Metabolism, and Intramyocellular and Hepatocellular Lipid Levels in Overweight Adults: A Randomized Clinical Trial. JAMA Netw Open 3: e2025454, 2020.

10. Rosato V, Edefonti V, Parpinel M, Milani GP, Mazzocchi A, Decarli A, Agostoni C, and Ferraroni M. Energy Contribution and Nutrient Composition of Breakfast and Their Relations to Overweight in Free-living Individuals: A Systematic Review. Adv Nutr 7: 455–465, 2016.

11. Pereira MA, Erickson E, McKee P, Schrankler K, Raatz SK, Lytle LA, and Pellegrini AD. Breakfast frequency and quality may affect glycemia and appetite in adults and children. J Nutr 141: 163–168, 2011.

12. Akbarzade Z, Mohammadpour S, Djafarian K, Clark CCT, Ghorbaninejad P, Mohtashami M, and Shab-Bidar S. Breakfast-Based Dietary Patterns and Obesity in Tehranian Adults. J Obes Metab Syndr 29: 222–232, 2020.

13. Iqbal K, Schwingshackl L, Gottschald M, Knüppel S, Stelmach-Mardas M, Aleksandrova K, and Boeing H. Breakfast quality and cardiometabolic risk profiles in an upper middle-aged German population. Eur J Clin Nutr 71: 1312–1320, 2017.

14. Chatelan A, Castetbon K, Pasquier J, Allemann C, Zuber A, Camenzind-Frey E, Zuberbuehler CA, and Bochud M. Association between breakfast composition and abdominal obesity in the Swiss adult population eating breakfast regularly. Int J Behav Nutr Phys Act 15: 115, 2018.

15. Maki KC, Phillips-Eakley AK, and Smith KN. The Effects of Breakfast Consumption and Composition on Metabolic Wellness with a Focus on Carbohydrate Metabolism. Adv Nutr 7: 613s–621s, 2016.

16. Clark CA, Gardiner J, McBurney MI, Anderson S, Weatherspoon LJ, Henry DN, and Hord NG. Effects of breakfast meal composition on second meal metabolic responses in adults with Type 2 diabetes mellitus. Eur J Clin Nutr 60: 1122–1129, 2006.

17. Wolever TM, and Miller JB. Sugars and blood glucose control. Am J Clin Nutr 62: 212S–221S; discussion 221S-227S, 1995.

18. Holst JJ, Gasbjerg LS, and Rosenkilde MM. The Role of Incretins on Insulin Function and Glucose Homeostasis. Endocrinology 162: 2021.

19. Holesh JE, Aslam S, and Martin A. Physiology, Carbohydrates. In: StatPearls. Treasure Island (FL): StatPearls Publishing Copyright © 2024, StatPearls Publishing LLC., 2024.

20. Hawari NS, Al-Shayji I, Wilson J, and Gill JM. Frequency of Breaks in Sedentary Time and Postprandial Metabolic Responses. Med Sci Sports Exerc 48: 2495–2502, 2016.

21. Jakubowicz D, Wainstein J, Landau Z, Raz I, Ahren B, Chapnik N, Ganz T, Menaged M, Barnea M, Bar-Dayan Y, and Froy O. Influences of Breakfast on Clock Gene Expression and Postprandial Glycemia in Healthy Individuals and Individuals With Diabetes: A Randomized Clinical Trial. Diabetes Care 40: 1573–1579, 2017.

22. Marathe CS, Rayner CK, Jones KL, and Horowitz M. Relationships between gastric emptying, postprandial glycemia, and incretin hormones. Diabetes Care 36: 1396–1405, 2013.

23. Crapo PA, Reaven G, and Olefsky J. Plasma glucose and insulin responses to orally administered simple and complex carbohydrates. Diabetes 25: 741–747, 1976.

24. Gannon MC, Nuttall FQ, Neil BJ, and Westphal SA. The insulin and glucose responses to meals of glucose plus various proteins in type II diabetic subjects. Metabolism 37: 1081–1088, 1988.

25. Polonsky KS, Given BD, Hirsch LJ, Tillil H, Shapiro ET, Beebe C, Frank BH, Galloway JA, and Van Cauter E. Abnormal patterns of insulin secretion in non-insulin-dependent diabetes mellitus. N Engl J Med 318: 1231–1239, 1988.

26. Franz MJ. Protein: metabolism and effect on blood glucose levels. Diabetes Educ 23: 643–646, 648, 650-641, 1997.

27. Bonuccelli S, Muscelli E, Gastaldelli A, Barsotti E, Astiarraga BD, Holst JJ, Mari A, and Ferrannini E. Improved tolerance to sequential glucose loading (Staub-Traugott effect): size and mechanisms. Am J Physiol Endocrinol Metab 297: E532–537, 2009.

28. Wajngot A, Grill V, Efendić S, and Cerasi E. The Staub-Traugott effect. Evidence for multifactorial regulation of a physiological function. Scand J Clin Lab Invest 42: 307–313, 1982.

29. Jovanovic A, Gerrard J, and Taylor R. The second-meal phenomenon in type 2 diabetes. Diabetes Care 32: 1199–1201, 2009.

30. Lee SH, Tura A, Mari A, Ko SH, Kwon HS, Song KH, Yoon KH, Lee KW, and Ahn YB. Potentiation of the early-phase insulin response by a prior meal contributes to the second-meal phenomenon in type 2 diabetes. Am J Physiol Endocrinol Metab 301: E984–990, 2011.

31. Lednev EM, Gavrilova AO, Vepkhvadze TF, Makhnovskii PA, Shestakova MV, and Popov DV. Disturbances in Dynamics of Glucose, Insulin, and C-Peptide in Blood after a Normalized Intake of a Mixed Meal in Obesity and Type 2 Diabetes Mellitus. Human Physiology 49: 668–674, 2023.

32. Moore MC, Smith MS, Farmer B, Coate KC, Kraft G, Shiota M, Williams PE, and Cherrington AD. Morning Hyperinsulinemia Primes the Liver for Glucose Uptake and Glycogen Storage Later in the Day. Diabetes 67: 1237–1245, 2018.

33. McCall AL. Insulin therapy and hypoglycemia. Endocrinol Metab Clin North Am 41: 57–87, 2012.

34. Mathew P, and Thoppil D. Hypoglycemia. In: StatPearls. Treasure Island (FL): StatPearls Publishing Copyright © 2024, StatPearls Publishing LLC., 2024.

35. Myers SR, McGuinness OP, Neal DW, and Cherrington AD. Intraportal glucose delivery alters the relationship between net hepatic glucose uptake and the insulin concentration. J Clin Invest 87: 930–939, 1991.

36. Dobbins RL, Davis SN, Neal DW, Cobelli C, Jaspan J, and Cherrington AD. Compartmental modeling of glucagon kinetics in the conscious dog. Metabolism 44: 452–459, 1995.

37. Moore MC, Smith MS, Farmer B, Kraft G, Shiota M, Williams PE, and Cherrington AD. Priming Effect of a Morning Meal on Hepatic Glucose Disposition Later in the Day. Diabetes 66: 1136–1145, 2017.

38. Galgani J, Aguirre C, and Díaz E. Acute effect of meal glycemic index and glycemic load on blood glucose and insulin responses in humans. Nutr J 5: 22, 2006.

39. Ahrén B, and Holst JJ. The cephalic insulin response to meal ingestion in humans is dependent on both cholinergic and noncholinergic mechanisms and is important for postprandial glycemia. Diabetes 50: 1030–1038, 2001.

40. Cherrington Adv. Banting Lecture 1997. Control of glucose uptake and release by the liver in vivo. Diabetes 48: 1198–1214, 1999.

41. Moore MC, Coate KC, Winnick JJ, An Z, and Cherrington AD. Regulation of hepatic glucose uptake and storage in vivo. Adv Nutr 3: 286–294, 2012.

42. Sindelar DK, Balcom JH, Chu CA, Neal DW, and Cherrington AD. A Comparison of the Effects of Selective Increases in Peripheral or Portal insulin on Hepatic Glucose Production in the Conscious Dog. Diabetes 45: 1594–1604, 1996.

43. Moore MC, Cherrington AD, Cline G, Pagliassotti MJ, Jones EM, Neal DW, Badet C, and Shulman GI. Sources of carbon for hepatic glycogen synthesis in the conscious dog. J Clin Invest 88: 578–587, 1991.

44. Hamilton KS, Gibbons FK, Bracy DP, Lacy DB, Cherrington AD, and Wasserman DH. Effect of prior exercise on the partitioning of an intestinal glucose load between splanchnic bed and skeletal muscle. J Clin Invest 98: 125–135, 1996.

45. Shah AM, and Wondisford FE. Tracking the carbons supplying gluconeogenesis. J Biol Chem 295: 14419–14429, 2020.

46. Edgerton DS, Cardin S, Emshwiller M, Neal D, Chandramouli V, Schumann WC, Landau BR, Rossetti L, and Cherrington AD. Small Increases in Insulin Inhibit Hepatic Glucose Production Solely Caused by an Effect on Glycogen Metabolism. Diabetes 50: 1872–1882, 2001.

47. Kuo T, McQueen A, Chen TC, and Wang JC. Regulation of Glucose Homeostasis by Glucocorticoids. Adv Exp Med Biol 872: 99–126, 2015.

48. Thau L GJ, Sharma S. Physiology, Cortisol. Treasure Island (FL): StatPearls Publishing.

49. Moore MC, Pagliassotti MJ, Swift LL, Asher J, Murrell J, Neal D, and Cherrington AD. Disposition of a mixed meal by the conscious dog. Am J Physiol 266: E666–675, 1994.

50. Hamman L, Hirschman II. Studies on blood sugar. Effect upon the blood sugar of the repeated ingestion of glucose. Johns Hopkins Hopsital Bulletin 344: 306–308, 1919.

51. Szabo AJ, Maier JJ, Szabo O, and Camerini-Davalos RA. Improved glucose disappearance following repeated glucose administration. Serum insulin growth hormone and free fatty acid levels during the Staub-Traugott effect. Diabetes 18: 232–237, 1969.

52. Abraira C, Buchanan B, and Hodges L. Modification of glycogen deposition by priming glucose loads: the second-meal phenomenon. Am J Clin Nutr 45: 952–957, 1987.

53. Jenkins DJ, Wolever TM, Taylor RH, Griffiths C, Krzeminska K, Lawrie JA, Bennett CM, Goff DV, Sarson DL, and Bloom SR. Slow release dietary carbohydrate improves second meal tolerance. Am J Clin Nutr 35: 1339–1346, 1982.

54. Gomis RR, Ferrer JC, and Guinovart JJ. Shared control of hepatic glycogen synthesis by glycogen synthase and glucokinase. Biochem J 351 Pt 3: 811–816, 2000.

55. Agius L. Role of glycogen phosphorylase in liver glycogen metabolism. Mol Aspects Med 46: 34–45, 2015.

56. Han H-S, Kang G, Kim JS, Choi BH, and Koo S-H. Regulation of glucose metabolism from a liver-centric perspective. Experimental & Molecular Medicine 48: e218–e218, 2016.

57. Agius L. Glucokinase and molecular aspects of liver glycogen metabolism. Biochemical Journal 414: 1–18, 2008.

58. Shiota C, Coffey J, Grimsby J, Grippo JF, and Magnuson MA. Nuclear import of hepatic glucokinase depends upon glucokinase regulatory protein, whereas export is due to a nuclear export signal sequence in glucokinase. J Biol Chem 274: 37125–37130, 1999.

59. Chu CA, Fujimoto Y, Igawa K, Grimsby J, Grippo JF, Magnuson MA, Cherrington AD, and Shiota M. Rapid translocation of hepatic glucokinase in response to intraduodenal glucose infusion and changes in plasma glucose and insulin in conscious rats. Am J Physiol Gastrointest Liver Physiol 286: G627–634, 2004.

60. von Wilamowitz-Moellendorff A, Hunter RW, García-Rocha M, Kang L, López-Soldado I, Lantier L, Patel K, Peggie MW, Martínez-Pons C, Voss M, Calbó J, Cohen PT, Wasserman DH, Guinovart JJ, and Sakamoto K. Glucose-6-phosphate-mediated activation of liver glycogen synthase plays a key role in hepatic glycogen synthesis. Diabetes 62: 4070–4082, 2013.

61. Roach PJ, Depaoli-Roach AA, Hurley TD, and Tagliabracci VS. Glycogen and its metabolism: some new developments and old themes. Biochem J 441: 763–787, 2012.

62. Aiston S, Green A, Mukhtar M, and Agius L. Glucose 6-phosphate causes translocation of phosphorylase in hepatocytes and inactivates the enzyme synergistically with glucose. Biochem J 377: 195–204, 2004.

63. Aiston S, Andersen B, and Agius L. Glucose 6-phosphate regulates hepatic glycogenolysis through inactivation of phosphorylase. Diabetes 52: 1333–1339, 2003.

64. Najjar SM, and Perdomo G. Hepatic Insulin Clearance: Mechanism and Physiology. Physiology 34: 198–215, 2019.

65. Rabkin R, Ryan MP, and Duckworth WC. The renal metabolism of insulin. Diabetologia 27: 351–357, 1984.

66. Stevenson RW, Cherrington AD, and Steiner KE. The relationship between plasma concentration and disappearance rate of immunoreactive insulin in the conscious dog. Horm Metab Res 17: 551–553, 1985.

