## supplement for "Duration of Morning Hyperinsulinemia Determines Hepatic Glucose Uptake and Glycogen Storage Later in the Day"

**Western Blotting**

**Sample Preparation and gel electrophoresis**

180 mg of frozen liver tissue was homogenized in 1 mL cold buffer (4 °C): 20 mM Tris, 200 mM NaCl, 50 mM NaF, 1mM EDTA, 1 mM EGTA, 10% glycerol (v:v), pH to 7.2. Protease inhibitors (#05892791001, MilliporeSigma, Burlington, MA) and phosphatase inhibitors (PHOSS-RO, MilliporeSigma) were added at the time of homogenization to prevent proteolysis and dephosphorylation of proteins. Homogenates rested on ice for 30 min and then were subjected to centrifugation (3000 rpm at 12 °C for 15 min). Supernatants were removed. Soluble protein content in each sample was quantified using the Bradford protein assay, where absorbance at 595 nm was measured using the SpectraMax M3 Multi-Mode Microplate Reader (Molecular Devices, Silicon Valley, CA). Samples were normalized to 2 ug/uL using homogenizing solution and 4x Laemmli sample buffer (#1610747, Bio-Rad, Hercules, CA). BSA was used for the standard curve (#23209, Thermo Fisher Scientific, Waltham, MA). Samples were boiled for 10 min and immediately cold-snapped in liquid nitrogen and stored at -80 °C. Aliquots containing 20 ug soluble protein were loaded onto SDS-polyacrylamide gels (#5671095, Bio-Rad) and then electrophoresed at 150V for ~ 90 min with running buffer (#1610732, Bio-Rad; 25 mM Tris, 192 mM glycine, 0.1% SDS, pH 8.3). A protein ladder was used for protein identification (#1610375, Bio-Rad), and laemmeli buffer was run in any lanes not containing a sample.

**Transfer and imaging**

### Proteins were semi-dry transferred to nitrocellulose membrane (P/N 926-31092, LI-COR BIOSCIENCES, Lincoln, NE) for 25 min at 15V using the Bio-Rad Trans-Blot SD Semi-Dry Transfer Cell apparatus (#1703848, Bio-Rad). Towbin transfer buffer (25 mM Tris, 192 mM glycine, 20% (v/v) methanol, pH 8.3: volume adjusted to 1L using dH2O) was used. Membranes were blocked using 5% BSA in TBST (1X TBST: 20 mM Tris; pH 7.5, 150 mM NaCl, 0.1% (w/v) Tween 20 detergent) for 1 hour at room temperature. Each membrane was incubated with the primary antibody at 4 °C overnight. Membranes were washed 3 x 10 min with TBST, incubated with appropriate HRP-conjugated secondary antibody for 1 hr, and washed 3 x 10 min with TBST again. 1 mL of HRP substrate (#WBKLS0500, MilliporeSigma) was added to each membrane. Membranes were immediately imaged using X-ray film (#Z370371, MilliporeSigma) and quantified using ImageJ software (http://imagej.nih.gov). Cyclophilin B was used as a loading control for all samples. Antibody dilutions were optimized for each protein. The following primary antibodies were used in western blotting analysis: pAkt (#9271, Cell Signaling Technology, Danvers, MA), tAkt (#9272, Cell Signaling Technology), pGS (#98348, Cell Signaling Technology), tGS (#3893, Cell Signaling Technology), pGSK3β (#9336, Cell Signaling Technology), tGSK3β (#9315, Cell Signaling Technology), and GK (#sc-17819, Santa Cruz Biotechnology). Anti-mouse and Anti-rabbit IgG (H+L), HRP conjugate secondary antibodies were purchased from Promega (Madison, WI).

**Real-time PCR**

25 mg of frozen canine liver tissue was homogenized using 600 μL of TRIzol reagent (#15596026, Thermo Fisher Scientific). RNA was purified using an RNA Mini-Prep kit (#R2050, Zymo Research, Orange, CA). The purity of RNA was verified based on A260/A280 ratios greater than 1.8. Total RNA was normalized to a final concentration of 5 μg. cDNA was synthesized using a cDNA reverse transcription kit (#43-688-14, Thermo Fisher Scientific) following the manufacturer’s instructions. This reaction was carried out using the MJ Mini Gradient Thermal Cycler (#PTC-1148, Bio-Rad). cDNA was stored at -20°C until use. Real-time PCR was performed using the CFX96 Dx System (Bio-Rad) with SYBR green fluorophore (#1725270, Bio-Rad; 10 μL SuperMix, 0.4 μM forward primer, 0.4 μM reverse primer, 0.2 μg cDNA template). Samples were run in duplicate. Primers were previously designed using OligoArchitect (MilliPoreSigma). Real-time PCR primer specificity was determined using BLAST analysis and melt curve analysis. The real-time PCR protocol included denaturation at 95°C for 3 min followed by 40 cycles of amplification (denaturation at 95°C for 10 s followed by annealing and extension at 55°C for 30 s). After each run, melt curve analysis was performed (65°C for 5 s followed by increments of 0.5°C every 5 s up to 95°C). GAPDH was used as the housekeeping gene for normalization and relative quantification using the 2^-ΔΔCt^ method (1).

**Supplemental Table 1. Nucleotide sequences for real-time PCR primers.**

| **Primer** | **Sequence (5’ → 3’)** | **Melting Temperature (T_m_)** |
| --- | --- | --- |
| GK - Forward | CAGAGGGGACTTTGAAATG | 59.7°C |
| GK - Reverse | CTGCATCTCCTCCATGTAG | 58.3°C |
| G6Pase - Forward | TGAAACTTTCAGCCACATCCG | 67.1°C |
| G6Pase - Reverse | GCAGGTAAAATCCAAGTGCGAA | 66.9°C |
| PEPCK - Forward | AGCTTTCAATGCCCGATTTCCAGG | 73.1°C |
| PEPCK - Reverse | TCAGCTCGATGCCGATCTTTGACA | 73.7°C |

**Glucokinase (GK) Activity Assay**

This assay was adapted from Dr. Masa Shiota (2). 60 mg of frozen canine liver tissue was homogenized in 1.5 mL of homogenization buffer (50 mM HEPES, 100 mM KCl, 5 mM MgCl_2_, 1 mM EDTA, 2.5 mM dithiothreitol (DTT), dH2O). Samples were subjected to ultracentrifugation at 30,000 rpm for 45 min at 4 degrees C (rotor type: TLA-55). The supernatant was isolated and kept on ice until use. NADH was used for the standard curve. 10 μL of each sample was combined with 190 μL of reaction medium (37°C: 2% albumin, 50 mM HEPES, 100 mM KCl, 7.5 mM MgCl_2_, 2.5 mM DTT, 0.5 mM NAD^+^, 5 mM ATP, 2 units of glucose-6-phosphate dehydrogenase). Samples were assessed at four different concentrations of glucose (0 mM, 0.5 mM, 7 mM, 100 mM). After the reaction medium and glucose were added, the plate was incubated at 37°C. The absorbance was read at 340 nm every 5 min for 30 min total. GK activity was determined at ½ V_max_ and V_max_.

**Glycogen**

Glycogen stores in frozen canine liver tissue were measured using the Keppler and Decker method (3). Briefly, 180 mg of tissue was homogenized with 0.6N perchloric acid (volume of perchloric acid added to each sample is equivalent to 5x the weight of tissue). 4, 5.5, and 7 mg of Oyster glycogen (#G8751, MilliPoreSigma) were homogenized in 1 mL perchloric acid and used as the three blanks. 200 μL of homogenate was combined with 100 μL KHCO_3_ (stock solution made by dissolving 1 g KHCO_3_ in 10 mL mpH_2_O) and 0.5 mL of amyloglucosidase-sodium acetate solution (#A7420, MilliPoreSigma: 2 mg/mL amyloglucosidase in sodium acetate buffer (0.4M: 19.8 g NaAc, 1L mpH_2_O, pH 4.8 adjusted using acetic acid)) was added to each sample. Samples were thoroughly mixed and placed in a 40 °C shaker bath for 2 hours. The glucose concentration in each sample was then measured using a glucose analyzer. Grams of glycogen per 100g of tissue was calculated and corrected using percent recovery determined by the oyster glycogen blanks. The average specific activity of glycogen was determined using liquid scintillation counting, which is utilized in the direct glycogen synthesis calculation.

**Glycogen Phosphorylase and Glycogen Synthase Activity Assay**

This assay was adapted from Dr. Masa Shiota (2). 200 μL of the reaction medium for glycogen phosphorylase (150 mM NaF, 1.5% bovine glycogen, 15 mM glucose-1-phosphate, 2 μCi [^14^C] glucose-1-phosphate/7 mL and 0.75 mM caffeine (for active phosphorylase) or 7.5 mM AMP (for active and inactive phosphorylase)) and glycogen synthase (50 mM Tris Buffer (pH 7.5), 5 mM EDTA, 1% bovine glycogen, 1.5 mM UDP-glucose, 2 μCi [^14^C] UDP-glucose/7 mL, and 15 mM Na_2_SO_4_ (for active synthase) or 3 mM glucose-6-phosphate (for active and inactive synthase)) was pre-incubated at 37 °C. 0.1 g of frozen liver canine tissue was homogenized with 1 mL of homogenization medium (50 mM Tris Buffer, 10 mM EDTA, 100 mM NaF, 5 mM DTT, 0.5% bovine glycogen). For the phosphorylase assay, a 1:10 dilution of the homogenate was made using the homogenization medium. A 1:5 dilution was made for the synthase assay. 100 μL of diluted homogenate was mixed well with the 200 μL pre-incubated reaction medium (37 °C). 50 μL of the incubated sample was spotted onto filter paper (20 mm x 25 mm). After the incubated medium was absorbed into the filter paper, the filter paper was placed in 70% ethanol (EtOH). While the samples remained incubating, this step was repeated after 20 min and again at 40 min. The filter papers were then washed in 70% EtOH, which was changed every 15 min for 60 min total. After a final 5 min wash with 100% ethanol, the filter papers were allowed to dry overnight. Each paper was placed in a scintillation vial and mixed for 2 hours in 1 mL DI water. 10 mL of Scintillation fluid was added to each sample, mixed, and kept for 24 hours. Radioactive counts were measured using liquid scintillation counting.

**Metabolite Analysis**

Blood samples were drawn and placed into tubes containing 1.6 mg/mL potassium EDTA. A 0.5 mL aliquot of blood from each sample tube was lysed with 1.5 mL of 4% perchloric acid and centrifuged. The supernatant was subsequently removed and stored at -80 °C until further use to quantitatively measure metabolites (lactate, alanine, and glycerol). 2 L stock solutions of buffer were prepared for each metabolite every 3 months (Suppl. Table 1). All buffers were dissolved in about 1500 mL mpH_2_O and brought up to 2 L after pH was adjusted.

| **Metabolite Being Measured** | **Reagents** | **Catalog Number (MilliporeSigma)** | **Amount Added** | **Additional Instructions** |
| --- | --- | --- | --- | --- |
| Lactate | Lactic acid (for std. curve) | L-2250 | 960 mg | Dissolved in 100 mL of mpH_2_O to achieve a final concentration of 100 mm/L |
|  | Glycine | G-7126 | 97.4 g | Adjust pH to 9.6 with ~28 g of buffer NaOH, then use 50% NaOH. Filter when done. |
|  | Hydrazine dihydrochloride | H-6628 | 53.2 g |  |
|  | Na_2_EDTA | ED2SS | 5.2 g |  |
| Alanine | Alanine (for std. curve) | A-7627 | 2.225 g | Dissolved in 100 mL of mpH_2_O to achieve a final concentration of 250 mm/L |
|  | Trizma base | T-1503 | 11.9 g | Add Na_2_EDTA after solids are completely dissolved. Adjust pH to 10 with 6.0 N HCl |
|  | Na_2_EDTA | ED2SS | 1.2 g |  |
|  | Hydrazine hydrate | H-0883 | 123 mL |  |
| Glycerol | Glycerol (for std. curve) | G-33 (Thermo Fisher) | 921 mg | Dissolved in 100 mL of mpH_2_O to achieve a final concentration of 100 mm/L |
|  | Glycine | G-7126 | 37.0 g | Adjust pH to 9.5 with 6.0 N HCl and 50% NaOH |
|  | MgCl_2_ – 6H_2_O | M33-600 | 5.0 g |  |
|  | Hydrazine hydrate | H-0883 | 123 mL |  |

Weekly stocks were made from the 3-month stocks by bringing 5.0 mL of lactate, 0.5 mL of alanine, and 0.38 mL glycerol up to a total volume of 50 mL with 3% perchloric acid each. These solutions were used for further analysis. 200 μL of each co-enzyme solution (lactate: 49 mL lactate buffer and 23.7 g NAD; alanine: 49 mL alanine buffer and 23.7 mg NAD; glycerol: 49 mL glycerol buffer, 87.4 g NAD, 87.4 g ATP, and 340.4 μL glycerol-3-phosphate dehydrogenase) was combined with 20 μL of each enzyme solution (lactate: 6 mL lactate buffer and 197 mL lactate dehydrogenase; alanine: 5.9 mL alanine buffer, 79 μL alanine dehydrogenase; glycerol: 5.8 mL glycerol buffer and 166 μL glycerokinase). Excitation was measured spectrophotometrically at 340 nm.

**Non-esterified Free Fatty Acids (NEFA) Analysis**

A 0.5 mL aliquot of plasma from each sample was stored in used in the assessment of NEFA’s. The Wako HR Series NEFA-HR (2) kit (Fujifilm, Tokyo, Japan) was utilized to quantitatively determine plasma NEFA levels. Absorbance was measured at 550 nm via enzymatic colorimetric methods following manufacturer instructions.

**References**

1.	**Livak KJ, and Schmittgen TD**. Analysis of relative gene expression data using real-time quantitative PCR and the 2(-Delta Delta C(T)) Method. *Methods* 25: 402-408, 2001.

2.	**Fujimoto Y, Torres TP, Donahue EP, and Shiota M**. Glucose Toxicity Is Responsible for the Development of Impaired Regulation of Endogenous Glucose Production and Hepatic Glucokinase in Zucker Diabetic Fatty Rats. *Diabetes* 55: 2479-2490, 2006.

3.	**Keppler D DK**. Glycogen determination with amyloglucosidase. *Methods of Enzymatic Analysis* 3: 1127-1131, 1974.
